# Adaptive transcriptional strategies underpin the host-specific virulence of the generalist oomycete *Phytophthora capsici* during early crown infection

**DOI:** 10.1101/2025.05.26.656185

**Authors:** Pablo Vargas-Mejía, Lino Sánchez Segura, Fernando U. Rojas-Rojas, Harumi Shimada-Beltrán, Julio C. Vega-Arreguín

**Author notes:** **Corresponding author:** JCVA.

## Abstract

*Phytophthora capsici* is a destructive, broad-host-range oomycete responsible for substantial losses in global agriculture. While most transcriptomic studies have focused on host responses, the mechanisms by which generalist pathogens dynamically adapt their infection programs to diverse plant species remain poorly understood. Here, we present a comparative transcriptomic analysis of *P. capsici* during early-stage crown infection in four taxonomically and immunologically distinct hosts, *Cucumis sativus, Cucumis melo, Capsicum annuum* (CM334), and *Solanum lycopersicum*, via RNA-seq and multiphoton microscopy. Focusing on crown infections, the natural entry point for the pathogen, we reveal host-specific transcriptional programs that underpin differential infection strategies and outcomes.

Our data show that *P. capsici* exhibits tightly regulated, host-dependent deployment of key virulence factors, including RxLR, NLP, and CRN, and elicitin effectors and reprograms its metabolism to exploit host-specific nutritional environments. In rapidly necrotizing hosts such as tomato, the pathogen induces glycolytic and fatty acid pathways while repressing immunogenic effectors. In contrast, cucurbits support prolonged biotrophic colonization, accompanied by the upregulation of carbohydrate metabolism and membrane transport genes. In the partially resistant chili pepper CM334, *P. capsici* shows signs of metabolic stress, cell wall remodeling, and effector repression, which is consistent with failed invasion.

Functional validation via RNAi-mediated silencing of selected effectors revealed distinct roles in modulating virulence and host necrosis, confirming the functional relevance of the transcriptomic profiles. Co-expression network analysis uncovered discrete transcriptional modules associated with tissue-specific colonization, nutrient acquisition, and immune evasion.

These results reveal how a generalist soil-borne pathogen finely tunes its gene expression in response to host-specific constraints, revealing conserved and host-specific transcriptional strategies that drive infection success or failure. This work provides mechanistic insight into adaptive virulence and expands our understanding of host-pathogen compatibility in eukaryotic microbes.

## 1 Background

*Phytophthora capsici* is a notorious filamentous phytopathogenic oomycete responsible for severe symptoms in vegetable, ornamental, and tropical crops, such as root and stem rot, reduced size in plants, and necrosis (Quesada-Ocampo et al. 2023). These symptoms lead to a yield decrease and, in severe cases, the loss of crops around the world. (Lamour et al. 2012b). As a soil-borne pathogen with a broad host range, *P. capsici* infects several species of the *Cucurbitaceae* and *Solanaceae* families, causing the first symptoms of diseases in the root, such as root rot and subsequent stem blight and fruit rot. *P. capsici* displays adaptative strategies to colonize its hosts, and it is considered a generalist phytopathogen; it has a highly adaptable infection strategy, allowing it to overcome diverse plant defense mechanisms (Quesada-Ocampo et al. 2023). Despite its importance, our understanding of the molecular mechanisms underlying host-specific infection strategies remains incomplete. Most of the previous studies have focused on plant transcriptomic responses to *P. capsici*, and fewer have investigated how the pathogen modulates its gene expression in response to different hosts and infection niches, providing valuable insights into pathogen adaptation and virulence (Chen et al. 2013; Jupe et al. 2013; Maillot et al. 2022). In addition, most transcriptomic studies of *P. capsici* have focused on foliar infection, even though this pathogen naturally enters plants mostly through soil and colonizes roots and the crown region (Hausbeck and Lamour 2004). The crown infection niche presents unique physiological and immunological features in terms of its structural characteristics, nutrient availability, and composition of host defense mechanisms that may influence pathogen gene expression and virulence strategies (Knight and Sutherland 2016; Stephens et al. 2008).

The early infection stage is particularly critical, as *P. capsici* must rapidly establish and grow within host tissues while evading recognition by host immune receptors. Pathogen success during this phase depends on the precise regulation of effector proteins, metabolic reprogramming, and stress adaptation mechanisms (Judelson 2017; Wang et al. 2023; Zuluaga et al. 2016). Notably, effector proteins such as RxLRs and Crinklers (CRNs) play essential roles in suppressing plant immune responses, whereas secreted hydrolytic enzymes facilitate tissue colonization (Chepsergon et al.; Stam et al. 2021; Wang et al. 2023; Zuluaga et al. 2016). On the other hand, metabolic plasticity also plays a key role, as *P. capsici* must exploit host-derived nutrients while resisting the oxidative stress imposed by plant defenses (Judelson 2017).

Despite these advances, the host-dependent transcriptomic responses of *P. capsici* remain poorly characterized, particularly in the context of crown infections in multiple hosts. Comparative transcriptomics of the pathogen across different plant species can reveal core virulence determinants as well as host-specific adaptations that shape infection outcomes. Understanding how *P. capsici* fine-tunes its gene expression in response to diverse host environments is crucial for developing targeted disease management strategies.

The aim of this study was to elucidate the transcriptomic responses of *P. capsici* during infection of four host species: two cucurbitaceous hosts (cucumber and melon) and two solanaceous hosts (CM334 chili pepper and tomato). Notably, our experimental design focused on infections initiated at the crown of the plant rather than the more commonly examined foliar infections. This approach is particularly relevant given that *P. capsici* is a soil-borne pathogen that exploits the crown region to establish systemic colonization. By investigating the transcriptomic profiles of *P. capsici* in these hosts at the early infection stage, we aimed to provide insights into the molecular determinants of host-specific virulence, as well as the metabolic adaptations that facilitate successful colonization.

## 2 Results

### 2.1 Host susceptibility and stem colonization

To evaluate host susceptibility to *P. capsici* D3, we infected the crowns of *C. annuum* (Criollo de Morelos 334 chili pepper), *C. melo* (melon), *C. sativus* (cucumber), and *S. lycopersicum* (tomato) plantlets with mycelia. The representative phenotypes of 5–8-week-old plants at 24 hours post-inoculation (hpi) are shown in Figure 1a-d. Among the tested hosts, cucumber exhibited the most pronounced aerial mycelial growth (Figure 1a-d), followed by melon and subsequently chili pepper and tomato. Interestingly, the extent of mycelial growth on the surface of the infected tissue contrasted with the time required for the pathogen to induce necrosis in the cucumber melon and tomato plants (Figure 1e).

**Figure 1:**
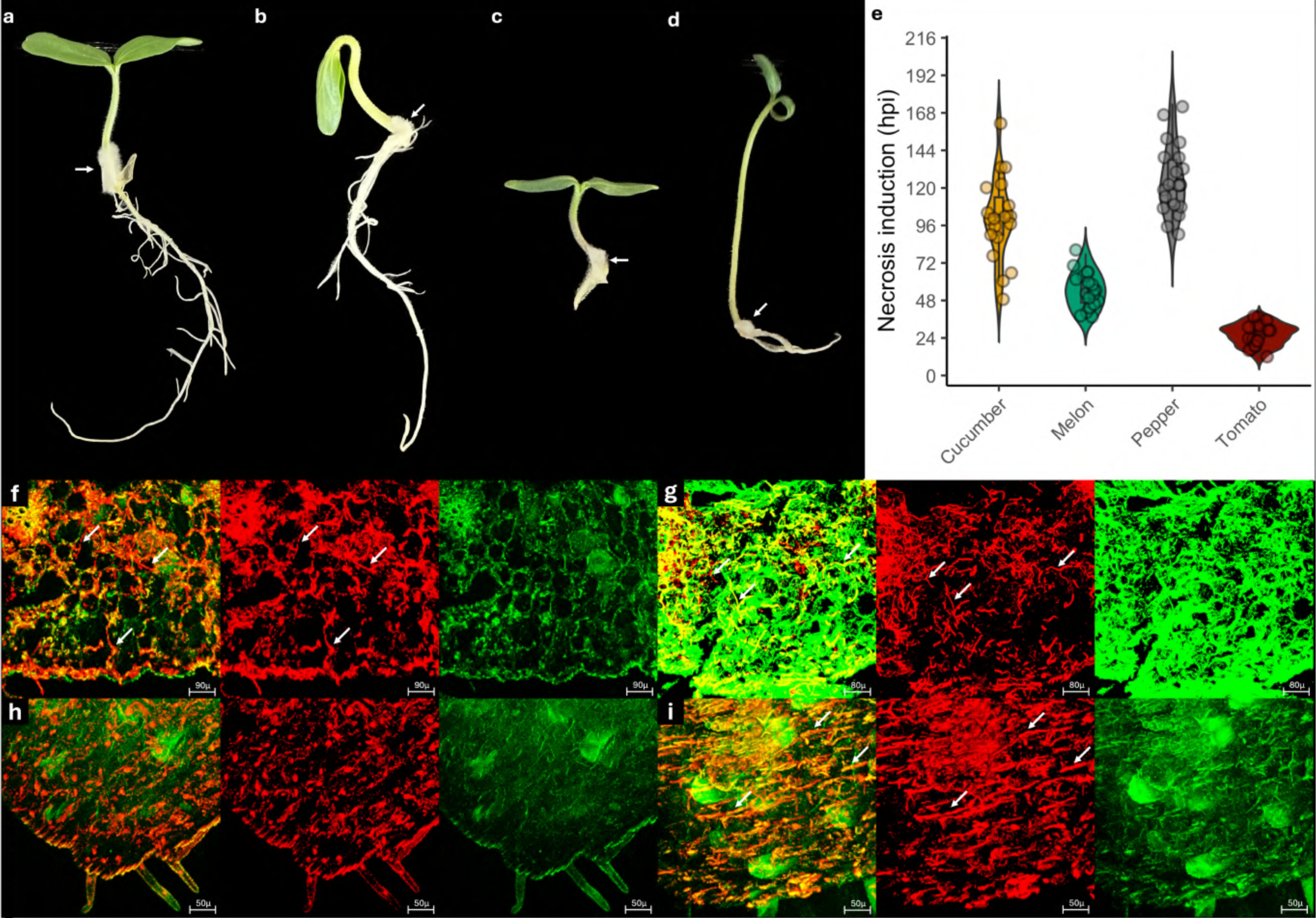
Dynamics of *P. capsici* infection. **a** Cucumber (24 hpi). **b** Melon (24 hpi). **c** Chili pepper (24 hpi). **d** Tomato (24 hpi). **e** Time to necrosis induction by the host. **f** 3D reconstruction of a stem section of a cucumber infected plant at 24 hpi. **g** 3D reconstruction of a stem section of a melon infected with *P. capsici* at 24 hpi. **h** 3D reconstruction of infected Chili pepper stem section (24 hpi). **i** 3D reconstruction of a stem section of a tomato infected with *P. capsici* (24 hpi). White arrows highlight portions of mycelium. The plant cell wall was stained with solophenyl flavine (green channel), and the hyphae of *P. capsici* were stained with propidium iodide (red channel). For each host, the left image presents both channels, the middle image presents only propidium iodine staining, and the right panel presents only the solophenyl channel.

The tomato was the most susceptible host; the first symptoms of necrosis were observed at 12 hpi for some individuals, and the median time for infected plants to show necrosis was 27 hpi, reflecting rapid progression of the infection. In contrast, the cucurbitaceous hosts displayed symptoms of necrosis at later times post-inoculation. Melon plants started to show necrosis at 36 hpi, with the median time of symptom appearance in the melon plant population at 54 hpi, whereas cucumber plants began to necrose at 48 hpi, reaching median necrosis onset at 102 hpi. Notably, chili pepper (CM334), a genotype previously reported to be partially resistant to *P. capsici* infection, exhibited an even further delayed necrotic response, with the first symptoms evident at 90 hpi and a median necrosis onset at 120 hpi (Figure 1e).

Despite delayed necrosis induction, in cucurbitaceous plants, greater intercellular mycelial growth was observed in these hosts during the sampling period. At 24 hpi, stem cross-sections dissected from 1 cm above the inoculation area showed abundant intercellular *P. capsici* D3 mycelial growth, particularly in vascular tissues, in an apoplastic manner (Figure 1f, g; Figure S1). In the tissue, the solophenyl (channel green) and propidium iodide (channel red) stains the cell walls of the plants and mycelia of *P. capsici* D3, respectively.

The tomato plants exhibited large mycelial growth. Unlike cucurbitaceous plants, the pathogen grows longitudinally into the surrounding parenchyma tissue but does not invade the vascular tissue (Figure 1i). Interestingly, the chili pepper plants did not exhibit intercellular mycelial growth in their stem tissues at 24 hpi (Figure 1h). Further examination of the inoculation site and 1 cm distal stems at 48 hpi confirmed that *P. capsici* D3 could not penetrate or invade chili pepper tissues effectively at these time points prior to necrosis (Figure S2).

### 2.2 Tissue-Specific Infection Dynamics: Root and Stem

To evaluate the tissue-specific pattern of infection, we examined intercellular mycelial growth in the roots of infected melon and chili pepper seedlings. At 24 hpi, *P. capsici* exhibited low proliferation in the roots of melon, and at 48 hpi, the pathogen showed a small increase in root colonization (Figure 2a). This could suggest that the pathogen had greater affinity for stems than for roots (Figure 1f-i and 2c-d).

**Figure 2:**
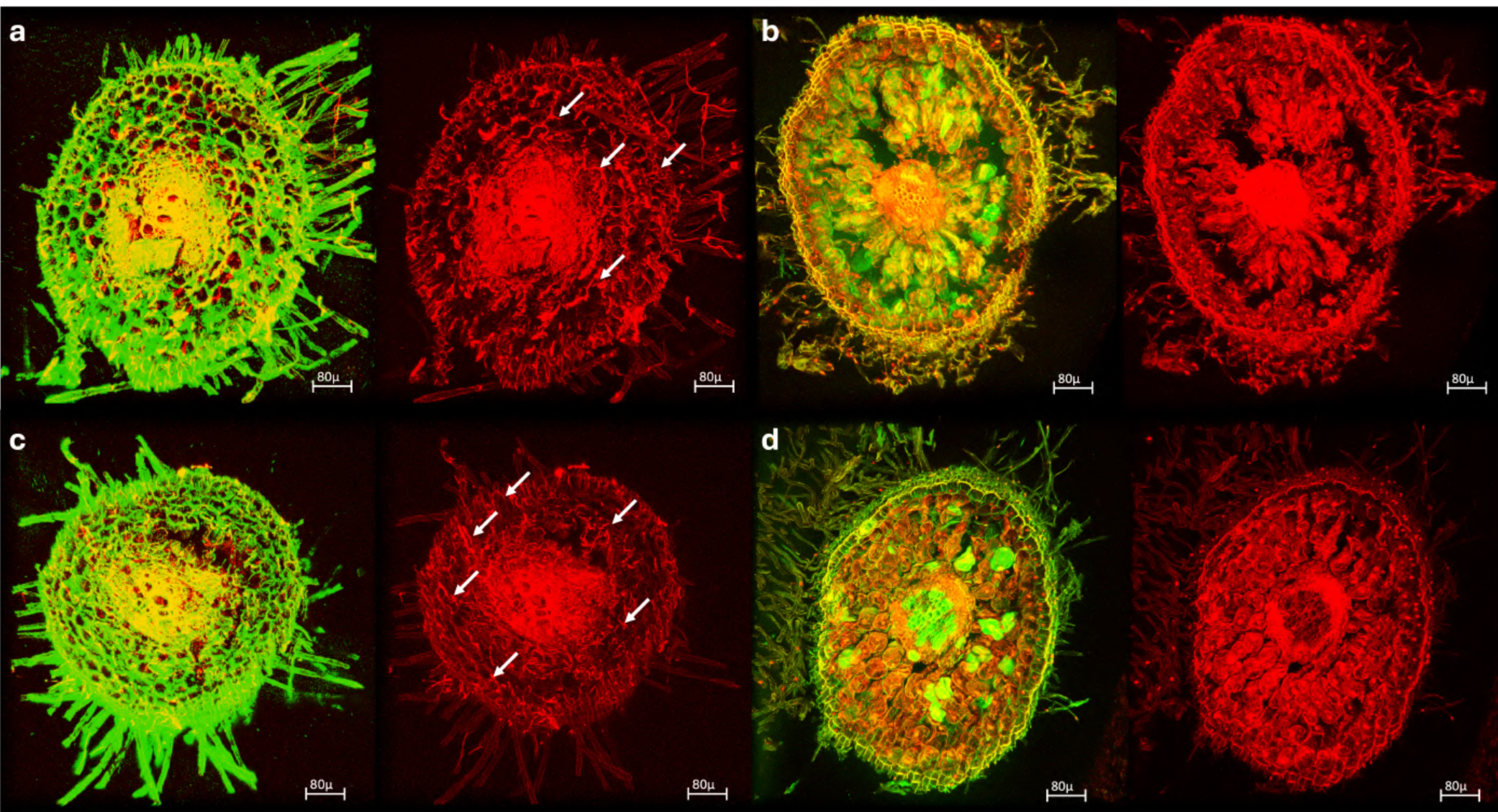
Root colonization of *P. capsici*. **a** 3D reconstruction of melon root at 24 hpi. **b** 3D reconstruction of chili pepper root at 24 hpi. **c** 3D reconstruction of melon root at 48 hpi. **d** 3D reconstruction of chili pepper root at 48 hpi. White arrows highlight portions of mycelium. The plant cell wall was stained with solophenyl flavine (green channel), and the hyphae of *P. capsici* were stained with propidium iodide (red channel). For each host, the left image presents both channels, and the right image presents only propidium iodine (PI) staining.

Consistent with observations at the stem, mycelial growth was not detected in chili pepper roots even when the pathogen developed appressoria at 24 and 48 hpi (Figure 2b and S3). At 96 hpi, infection progression resulted in *P. capsici* mycelial growth extending into the aerial photosynthetic parts of tomato; at 120 hpi in melon and cucumber; and at 144hpi in pepper chili plantlets. This observation suggests that *P. capsici* tends to colonize the aerial portions of its hosts, despite initial zoospore deposition being concentrated at the crown, roots, and stem base under field conditions (Figure S4).

### 2.3 Comparative transcriptomic analysis reveals host-dependent pathogenic strategies

We decided to perform RNA-Seq of *P. capsici* infecting Solanaceous and Cucurbitaceous hosts to understand the transcriptional response of the pathogen to the diverse environments that each host represents. Mapping to the reference transcriptome yielded an average mapping rate of 77% for each library. After principal component analysis, we found an evident separation of the transcriptome responses of the *Phytophthora* samples when the pathogen was infecting a host compared with the pathogen growing in V8 agar media (Figure 3a). Additionally, PCA showed an aggrupation of the transcriptomic responses of the cucurbits and tomato infections, although 1 tomato sample was grouped with the chili pepper samples (Figure 3a).

**Figure 3:**
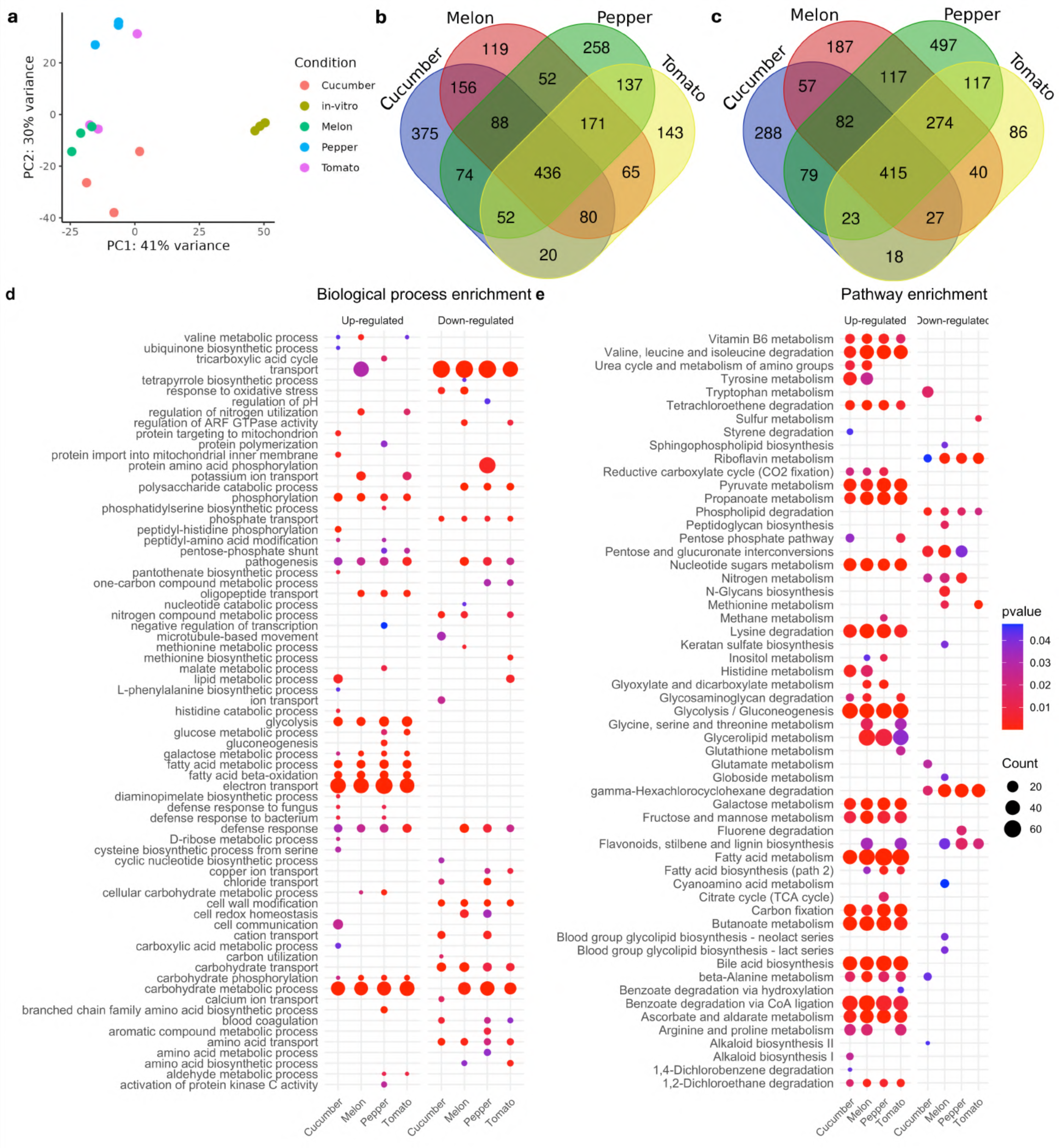
Transcriptional responses of *P. capsici* to different hosts. **a** Principal component analysis of *P. capsici* transcript expression. **b** Venn diagram of up-regulated DEGs. **c** Venn diagram of down-regulated DEGs. **d** Biological process and **e** KEGG pathway enrichment of *P. capsici* differentially expressed genes.

Among the 19805 transcripts of *P. capsici*, 4470 genes were found to be differentially expressed (DEGs = padj < 0.05 and Log2FC > 1 OR < -1) among all the conditions compared with our control pathogen growing in V8 media (Table 1). We found, on average, approximately 2300 DEGs for each sample, with tomato infections resulting in fewer differentially expressed genes and chili pepper infections resulting in the most differentially expressed genes (Table 1). Among the 4470 DEGs, only 436 were induced among all the conditions, and 415 were down-regulated, indicating that the core transcriptome of *P. capsici* infecting four different hosts is not composed of a large number of genes (Figure 3b, c). This core set of DEGs shared across all *P. capsici* infection conditions was subjected to Gene Ontology (GO) and Kyoto Encyclopedia of Genes and Genomes (KEGG) pathway enrichment analyses to identify key biological processes and pathways involved in plant-pathogen interactions. Among the biological processes, significant enrichment was observed for processes associated with metabolic activity, including fatty acid metabolism, carbohydrate metabolic processes, and glycolysis, alongside defense-related processes such as pathogenesis and electron transport (Figure S5a). Additionally, terms related to cell wall modification and phosphate transport were prominent, reflecting infection and nutrient acquisition by the pathogen.

**Table 1:** Differentially expressed genes*.

| Condition | Total genes | Up regulated | Down regulated |
| --- | --- | --- | --- |
| Cucumber | 2270 | 1281 | 989 |
| Melon | 2366 | 1167 | 1199 |
| Chili pepper | 2872 | 1268 | 1604 |
| Tomato | 2104 | 1104 | 1000 |
| Total Unique | 4470 | 2226 | 2307 |

\* $p_{adj} < 0.05$ $\text{Log}_2\text{FC} > 1$ OR $< -1$

Pathway enrichment analysis revealed a diverse range of metabolic pathways (Figure S5b). These included fatty acid beta-oxidation, glycolysis/gluconeogenesis, and amino acid metabolism, suggesting a reconfiguration of primary metabolic networks during infection. Notably, pathways associated with the mobilization of nutrients from the host, such as glycerolipid and galactose metabolism, were upregulated, emphasizing the pathogen’s strategy to exploit host resources. In parallel, the enrichment of pathways related to defense mechanisms, including alkaloid biosynthesis and gamma-hexachlorocyclohexane degradation, underscores the activation of plant immune responses. The differential enrichment of these pathways among the upregulated and downregulated DEGs provides insights into the dynamic interplay between metabolic reprogramming and adaptation to host defense during *P. capsici* infection.

The transcriptional response of *P. capsici* revealed remarkable adaptability to the unique micro ecological niches generated by its hosts. While some processes are shared, the pathogen’s specific enrichment varies considerably, reflecting its ability to exploit host-specific vulnerabilities and overcome distinct physiological and defense barriers.

In solanaceous hosts, particularly tomato, *P. capsici* prioritized pathways associated with rapid energy acquisition, such as glycolysis and fatty acid metabolism. This metabolic change was probably due to the faster strategy of pathogen colonization, which corresponds to the host’s early necrotic response. Interestingly, during tomato infection, *P. capsici* is also enriched in alkaloid degradation pathways, suggesting that the pathogen may counteract host-produced antimicrobial compounds, including alkaloids, to establish infection. The capacity of *P. capsici* to degrade these compounds highlights the intense chemical warfare between tomato and *P. capsici*, with the pathogen leveraging its metabolic flexibility to neutralize host defenses (Figure 3d, e).

In contrast, infection of the partially resistant chili pepper CM334 resulted in a distinct pattern of transcriptional enrichment in the pathogen (Figure 3). Processes related to cell wall modification and lipid metabolism were notably upregulated in *P. capsici* during its attempted colonization of chili pepper. These enrichments, including pathways such as terpenoid and glycerolipid metabolism, suggest that *P. capsici* was preparing to breach structural barriers and colonize the host. However, the absence of detectable pathogen growth within chili pepper tissues at 24 hpi (Figure 1c and h) implies that the mechanism of infection was insufficient to overcome the physical and biochemical defenses of this plant. Additionally, the enrichment of nitrogen compound metabolism pathways in *P. capsici* during chili pepper infection may indicate an effort to adapt to nutrient scarcity, reflecting a limitation imposed by host resistance mechanisms.

Melon infection was characterized by strong enrichment in carbohydrate metabolism, including the galactose and glycerolipid pathways, and processes such as nitrogen utilization and protein glycosylation (Figure 3d, e). These adaptations demonstrate the capacity of the pathogen to use the host’s abundant resources to support its growth and effector production. Similarly, in cucumber, *P. capsici* was significantly enriched in glycolysis and gluconeogenesis, suggesting an intensified focus on extracting energy and carbon from the host. Pathways related to transport, such as phosphate and copper transport, were also highly enriched in cucurbitaceous hosts. In chili pepper, the pathogen appears to invest heavily in preparatory mechanisms to overcome structural and chemical defenses, albeit unsuccessful in this case. Conversely, in more susceptible hosts, *P. capsici* capitalizes on the availability of nutrients and the relative lack of effective defensive barriers to drive its infection forward.

### 2.4 Host-dependent effector expression profiles

We found that 70 RxLRs, 28 Elicitins, 18 CRNs, and 17 NLPs effectors were differentially expressed. RxLR expression was mainly down-regulated except for 12 genes, and its expression was grouped according to the family of the infected host; the same behavior was found for the CRN effectors (Figure 4). However, Elicitins clustered with melon, tomato, and chili pepper, with the cucumber condition as the most different. NLPs expression was clustered for tomato, cucumber, and melon, with chili pepper being the most dissimilar condition for the expression of this effector family. NLPs were also repressed, with only 5 NLPs being overexpressed.

**Figure 4:**
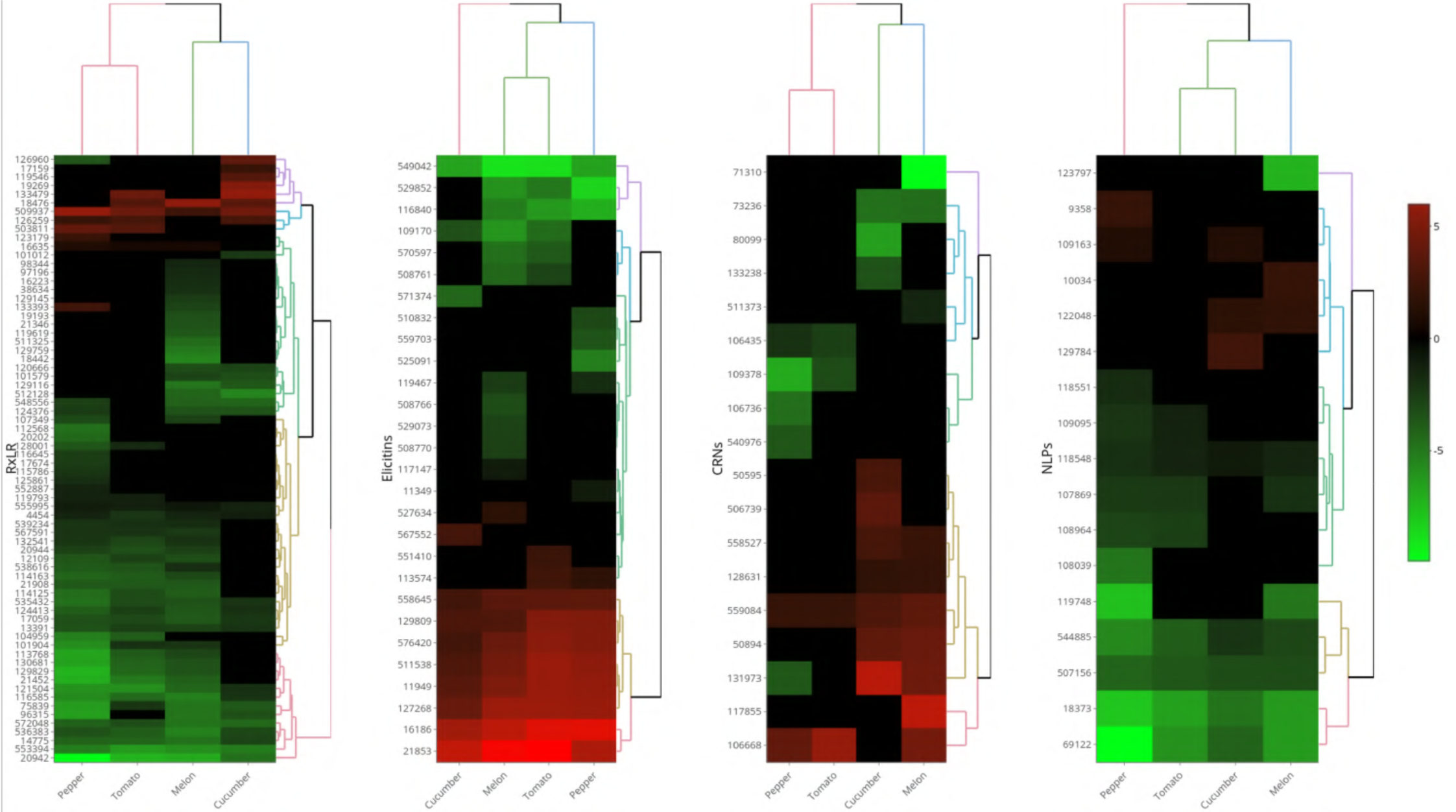
Heatmap of the Log2FC values of differentially expressed *Phytophthora* RxLR, elicitin, CRN, and NLP effectors.

A review of the annotated RxLR effectors in our dataset revealed a diverse arsenal with distinct expression patterns that align with previously reported functions. For example, several homologs of PsAvh328 (genes Pc123179, Pc128001, and Pc552887) have been shown in earlier studies to interfere with host defense signaling and modulate immune responses (Wang et al. 2011). Interestingly, while some Avh328 homologs remain unexpressed under some conditions, gene Pc123179 is induced in chili pepper, suggesting a specialized role in overcoming host-specific defenses. Similarly, PsAvh16 (Pc14944, Pc14948, and Pc533084) has been implicated in suppressing programmed cell death, thereby assisting the pathogen in establishing a biotrophic interaction (Fedkenheuer 2016). Gene Pc13391, which encodes the uncharacterized Avh166 gene, is consistently downregulated across all hosts, possibly indicating its involvement at earlier/later infection stages or under environmental cues not captured in our current sampling. The marked downregulation of Pc20942 across conditions may reflect tightly regulated deployment. The effector PcAvh103 (gene Pc536383) has been linked to the manipulation of EDS1, thus facilitating virulence; however, our data revealed that this gene was downregulated in all host infections (Li et al. 2020).

Our expression analysis of elicitin-encoding genes in *P*. *capsici* revealed notable variability across host conditions. Several elicitins, including the genes Pc16186, Pc21853, and Pc127268, exhibit robust positive expression in all hosts. However, the gene Pc508761 (annotated as INF2B) is strongly repressed in melon and tomato, and the gene Pc508770 (INF1-like) is not induced in cucumber, chili pepper, or tomato but is moderately repressed in melon. INF1 is one of the best-characterized elicitins and is known in *P. infestans* for its sterol-binding activity and ability to elicit a hypersensitive response in a range of plant hosts (Ah-Fong et al. 2021; Du et al. 2017). Additionally, INF2B is related to triggering HR (Huitema et al. 2005). The observed repression of both INF1 and INF2B in melon and INF2B in tomato suggest that *P. capsici* may actively downregulate these elicitors to avoid triggering host defense responses and avoid necrosis during its biotrophic stage.

Crinkler (CRN) effectors form a large and diverse family in oomycetes that can manipulate host cell death and immune responses. In our dataset, the CRN125 homolog (gene Pc131973) is strongly expressed in cucumber and melon but is downregulated in chili pepper and not in tomato, suggesting a role tailored to the cucurbit host environment, as proposed to interact with cucurbit TCP14-1 or TCP14-2 to enhance virulence (Stam et al. 2021). Multiple entries annotated as CRN15 show variable patterns; for example, the Pc50894 gene is expressed in both cucumber and melon but not in other hosts, whereas the Pc106668 also annotated as CRN15, is upregulated in melon, chili pepper, and tomato. In contrast, CRN10 (gene Pc506739) is expressed solely in cucumber, and CRN10 homologs (genes Pc109378 and Pc80099) are downregulated in chili pepper and tomato, suggesting distinct regulatory controls even among closely related CRNs. Other CRN annotations, such as CRN77 (gene Pc128631) and CRN34 (gene Pc117855), exhibit modest expression in specific hosts (CRN77 in cucumber and melon and CRN34 prominently in melon), whereas members such as CRN5 (gene Pc540976) and CRN21 (gene Pc133238) show little to no differential expression, possibly reflecting functions that are restricted to particular infection stages or conditions. Pc559084 was induced in all hosts, and this gene was shown to be able to suppress HR triggered by INF1 but increase disease susceptibility in *N. benthamiana* plants (Chen et al. 2015). Many CRNs in *P. capsici* remain uncharacterized. The diverse expression profiles observed here underscore the potential for functional specialization within the CRN family and highlight the need for further research to unravel their roles in host-specific pathogenicity.

Nep1-like proteins (NLPs) are a diverse family of secreted effectors known to induce necrosis in dicotyledonous plants by binding to membrane glycosylinositol phosphorylceramides (GIPCs) (Lenarčič et al. 2017). In our dataset, the expression profiles of several NLPs revealed distinct, host-dependent regulation. For example, the gene Pc9358, annotated as PcNLP14, is uniquely induced in chili pepper, whereas its expression does not change in cucumber, melon, or tomato. PcNLP14 was previously shown to be one of the most highly expressed NLP effectors in chili pepper and *N. benthamiana,* and its agroinfiltration in plants resulted in the induction of necrosis (Feng et al. 2014). In contrast, the Pc122048 gene was moderately induced in both cucumber and melon but was not differentially expressed in chili pepper and tomato. Pc129784 PcNLP2, another inducer of necrosis, is exclusively induced in cucumber, further supporting host-specific functions among NLPs (Chen et al. 2018). On the other hand, several NLPs, including the gene Pc107869 and its close homolog gene Pc108964, are consistently repressed in melon, chili pepper, and tomato, indicating that their expression could be tightly controlled to avoid premature host cell death (Chen et al. 2018). Other NLPs, such as Pc108039 NLP4 and Pc119748 NLP6, exhibit strong repression specifically in chili pepper and melon, whereas the genes Pc18373 and Pc69122 are downregulated across all hosts.

### 2.5 Host- and Tissue-Specific Phenotypes Associated with Effector Gene Silencing

We investigated the role of four *P. capsici* genes by silencing via direct incubation with dsRNA. The targeted genes included the differentially expressed necrosis- and ethylene-inducing peptide-like protein (NLP Pc9358), induced exclusively in chili pepper; the RxLR effector Pc503811, induced in chili pepper and tomato; the RxLR effector Pc18476, induced across all susceptible hosts (cucumber, melon, and tomato) (Figure 4); and a non-differentially expressed conserved cell cycle gene (Pc533030, encoding SDA1), known to be essential for sporangial morphology, mycelial growth, and virulence (Zhu et al. 2016). Pathogen mycelia were incubated with the corresponding dsRNAs for 2 h prior to infection assays on detached leaves and plantlet stems near the crown to better mimic the conditions from previous experiments.

After 96 hpi, the control *P. capsici* grew on detached cucumber leaves. In contrast, all the following dsRNA treatments decreased pathogen growth: Pc503811 (18.24%), Pc9358 (65.29%), PcSDA1 (76.47%), and Pc18476 (78.82%) (Figure 5a, e). For detached tomato leaves measured at 72 hpi, PcSDA1 dsRNA treatment unexpectedly resulted in a 10.58% increase in pathogen growth, whereas the Pc503811 and Pc9358 treatments reduced growth by 76.51% and 84.6%, respectively (Figure 5b, c, e).

**Figure 5:**
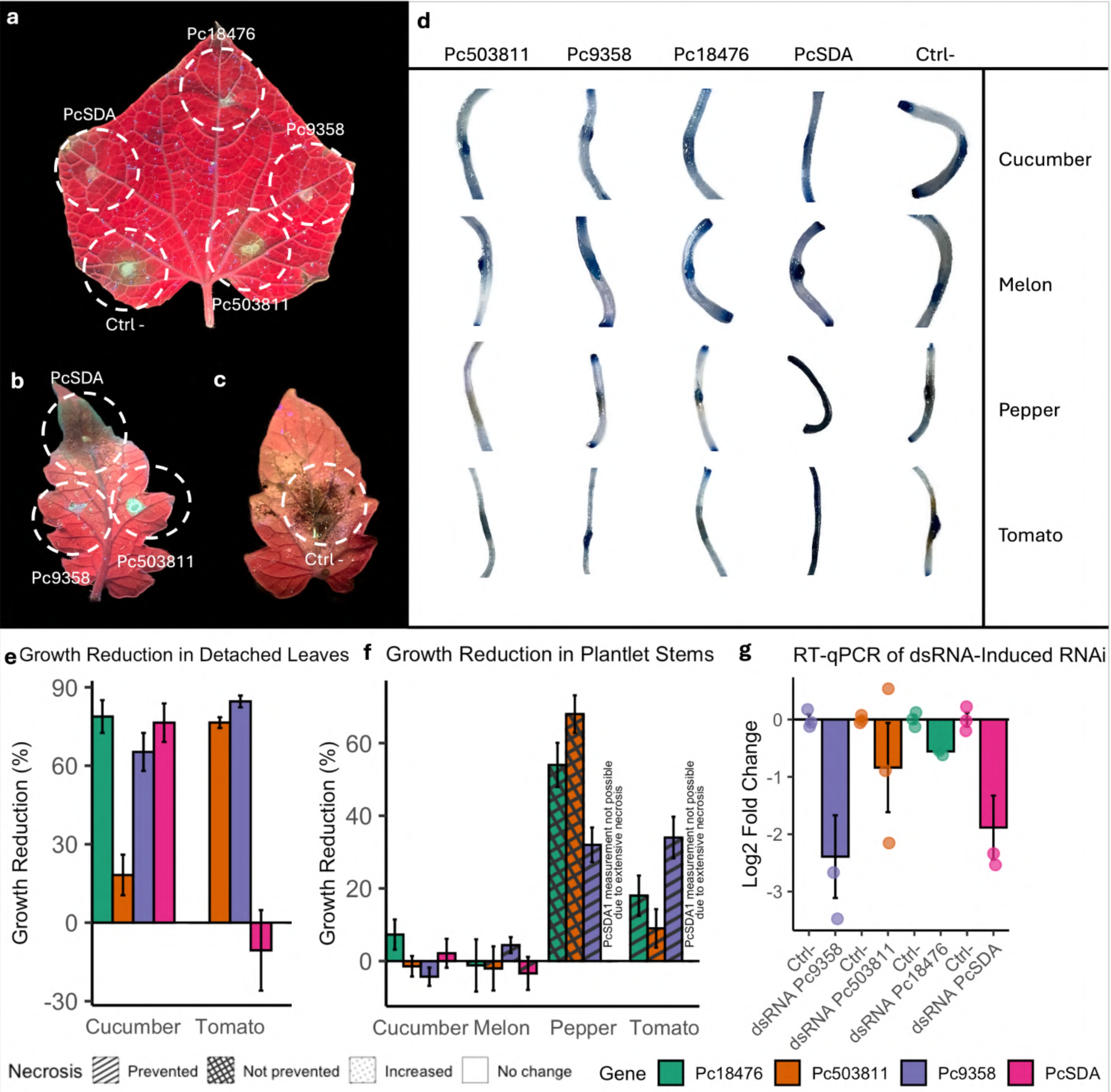
*P. capsici* gene-silenced phenotypes. **a** Cucumber detached leaf inoculated with *P. capsici* treated with dsRNAs targeting PcSDA, Pc18476, Pc9358 and Pc503811 or control plants treated with dsRNA buffer without dsRNA production after 96 hpi. **b** Tomato leaf inoculated with dsRNA-treated *P. capsici* after 72 hpi. **c** Tomato leaf inoculated with control *P. capsici* after 72 hpi. **d** Trypan blue-stained stems of cucumber, melon, tomato and chili pepper plantlets inoculated with control or dsRNA-treated *P. capsici* after 96 hpi. **e** Growth reduction measurement of detached leaves infected with dsRNA-treated *P. capsici*. **f** Growth reduction measurement of plantlet stems infected with dsRNA-treated *P. capsici*.

In plantlet stem infection (96 hpi), cucumber did not differ between the dsRNA treatment groups and the control group (Figure 5d, f). Similarly, in melon stems infected with dsRNA-treated pathogen exhibited growth comparable to that of the control, although dsRNA treatments notably prevented the appearance of necrosis. Conversely, in chili pepper and tomato stems, the PcSDA1 dsRNA-treated pathogen induced aggressive necrosis, surpassing the control severity and causing complete tissue necrosis. In chili pepper, the Pc18476 and Pc503811 dsRNA treatments resulted in reduced pathogen growth by 54% and 68%, respectively, but failed to prevent necrosis. Notably, while the Pc9358 dsRNA treatment caused a more moderate 32% growth reduction, it completely prevented necrosis in the chili pepper stems (Figure 5d, f). In tomato stems, dsRNA treatments targeting Pc18476, Pc9358, and Pc503811 effectively prevented necrosis and resulted in pathogen growth reductions of 18%, 34%, and 9%, respectively, relative to the control. Corroboration of gene silencing induction by dsRNA treatment was performed via RT-qPCR (Figure 5g).

### 2.6 Co-expression analysis uncovers distinct transcriptional modules

The co-expression analysis of *P. capsici* during infection of cucumber, melon, chili pepper, and tomato revealed 10 distinct gene clusters with host-specific expression patterns and functional enrichments. Cluster 1 (1,999 genes) is expressed at low levels in cucumber but is strongly induced in melon, tomato, and chili pepper; it is enriched in processes such as ubiquitin-dependent protein catabolism, vesicle-mediated transport and protein complex assembly. These functions, along with pathways such as propanoate and fatty acid metabolism as well as secretion components, suggest that in hosts other than cucumber, the pathogen deploys rapid protein turnover and secretion mechanisms to modulate its virulence arsenal effectively (Figure 6).

**Figure 6:**
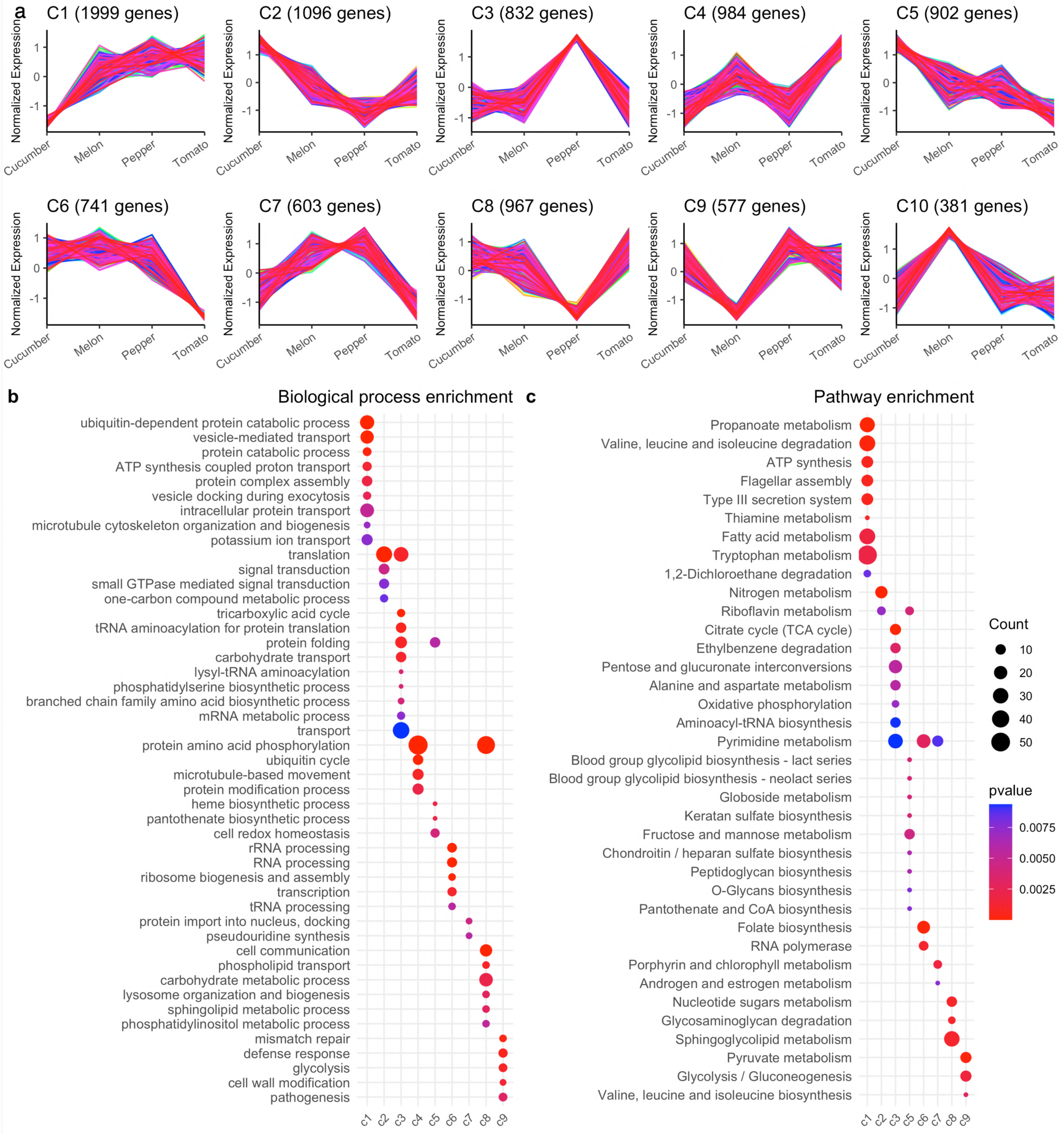
Clustering of co-expressed genes based on their expression profiles across infected hosts. **a** Each panel represents a cluster (C1-C10) with the number of genes indicated in parentheses. **b** Biological process and **c** KEGG pathway enrichment of *P. capsici* cluster genes.

Cluster 2 (1,096 genes) was mostly active in cucumber, with intermediate levels in melon and tomato and low expression in chili pepper. The enrichment of translation and nitrogen metabolism within this cluster implies a heightened emphasis on protein synthesis and energy acquisition during the more tolerant, delayed infection observed in cucumber. Cluster 3 (832 genes) was specifically upregulated in chili pepper and strongly enriched in central metabolic pathways, including the tricarboxylic acid cycle, oxidative phosphorylation, and carbohydrate transport, which may reflect an attempt by *P. capsici* to overcome the robust defense responses characteristic of this partially resistant host.

A distinct transcriptional signature is observed during tomato infection, where cluster 4 of 984 genes is predominantly induced. The overrepresentation of genes involved in protein amino acid phosphorylation, the ubiquitin cycle, and microtubule-based movement in this group suggests that rapid cytoskeletal reorganization and signal transduction are critical for the early, aggressive colonization that leads to swift necrosis in tomato. Moreover, cluster 5, with 902 genes highly expressed in cucumber at intermediate levels in melon and chili pepper but low in tomato, was enriched in pathways related to redox homeostasis, heme and pantothenate biosynthesis, and glycolipid metabolism. This profile indicates that in cucumber, the lipid biosynthetic and redox-balancing processes of the pathogen may be fine-tuned to support prolonged infection.

Cluster 6 with 741 genes upregulated in cucumber, melon, and chili pepper but repressed in tomato; its enrichment in rRNA processing, ribosome biogenesis, transcription, and tRNA processing underscores the importance of enhancing the pathogen’s translational machinery in hosts where colonization occurs over an extended period. Cluster 7 (603 genes), with moderate expression in cucumber and high expression in melon and chili pepper, was associated with the modulation of pathways such as porphyrin and pyrimidine metabolism, suggesting a role in fine-tuning gene expression and nuclear-cytoplasmic transport in response to host defenses.

Notably, a large set of 967 genes in cluster 8 was highly expressed in cucumber, melon, and tomato but was downregulated in chili pepper. This group was enriched for processes involved in cell communication, protein phosphorylation, phospholipid transport, and carbohydrate metabolism, along with nucleotide sugar and sphingoglycolipid metabolism. Such enrichment may reflect the active remodeling of membranes and signaling networks to optimize effector secretion and host interactions in more susceptible hosts. In addition, cluster 9, comprising 577 genes (upregulated in cucumber, chili pepper, and tomato with low expression in melon), was enriched in pathways including mismatch repair, defense response, glycolysis, cell wall modification, and pathogenesis (Figure 6). Finally, a smaller, melon-specific cluster 10 of 381 genes, despite yielding no robust enrichment.

## 3 Discussion

### 3.1 Host-specific susceptibility and pathogen adaptation

*Phytophthora capsici* is a highly adaptable oomycete pathogen with a broad host range. This wide host range is facilitated by its ability to infect both root and aerial tissues, employ diverse infection strategies, and rapidly evolve virulence traits. However, the extent and nature of susceptibility vary markedly among host species and are influenced by both preformed defenses and inducible immune responses (Quesada-Ocampo et al. 2023). Our study demonstrated that host susceptibility to *P. capsici* is not a uniform phenomenon but is distinctly modulated by host species. In tomato plants, a Solanaceous host, necrosis is visible as early as 12 hours post-inoculation for some individuals, with a population median onset of 27 hpi, suggesting rapid cell death, which allows further pathogen colonization (Singh and Upadhyay 2014). In contrast, cucurbitaceous hosts (cucumber and melon) exhibit delayed necrosis (first symptoms at 36-48 hpi with medians ranging from 54 to 102 hpi), yet these hosts support extensive intracellular colonization. This combination of delayed cell death and significant intracellular proliferation may reflect a more tolerant interaction. As defined in plant pathology, tolerance refers to a plant’s ability to endure the presence of a pathogen without suffering significant loss in yield or quality, even though the pathogen may still infect and reproduce within the plant (Pagán and García-Arenal 2018).

Our *Phytophthora* propagation findings in cucumber and melon align with previous studies that have documented a prolonged biotrophic phase in certain *Phytophthora* interactions. For example, in *P. infestans* infections of potato, delayed necrotic induction is associated with sustained colonization, which allows the pathogen to build a critical biomass before a switch to necrotrophy (Lee and Rose 2010; Schmelzer et al. 1995). In our system, the extensive vascular colonization observed in cucumber and melon via multiphoton microscopy suggests that these hosts may provide a relatively permissive environment that limits early defense activation. The result is a biotrophic scenario in which *P. capsici* can establish a systemic infection while minimizing the rapid release of damage-associated molecular patterns (DAMPs), which typically trigger strong immune responses (Hernández-Chávez et al. 2017). Conversely, chili pepper (CM334), which was previously characterized as partially resistant (López-Martínez et al. 2011), shows delayed necrosis coupled with an absence of intracellular colonization. We indeed observed that *P. capsici* was unable to penetrate chili pepper plants. These findings indicate that in resistant hosts, preformed or induced defenses, such as reinforced cell walls and rapid antimicrobial signaling, effectively restrict pathogen entry and subsequent proliferation.

Other studies have documented similar patterns. For example, (Maillot et al. 2022) compared the transcriptomes of *P. capsici* strains infecting partially resistant CM334 chili peppers with those infecting susceptible cultivars and reported that, in resistant tissues, the proliferation of the pathogen was blocked early by rapid defense signaling. In contrast, in susceptible tissues, the pathogen has the capacity to change the transcriptomic programming of the host response to delay necrosis and establish more robust colonization. Our findings extend these observations to cucurbits, suggesting that the apparent tolerance of cucumber and melon could allow pathogen proliferation by a more extended period for nutrient use and colonization.

### 3.2 Unique aspects of crown infection and broader implications

A novel focus of our study was infection initiation at the stem base and root crown rather than in foliar tissues. Most studies on *Phytophthora* have focused on leaf infections (Chen et al. 2018; Li et al. 2020; Zhu et al. 2016). However, pathogens such as *P. capsici* are transmitted by the soil and naturally target the crown region where roots and stem bases interface with the soil environment. By examining these infection areas, we have uncovered critical insights into the early stages of pathogen establishment. The crown region represents a crucial area with physical barriers and localized immune responses that determine the success of infection (Bostock et al. 2014; Piccini et al. 2019). In fact, the results here revealed that *P. capsici* can establish extensive intracellular colonization in the crown of cucurbitaceous hosts without triggering rapid necrosis, suggesting that these tissues may offer a relatively protected niche that permits sustained pathogen growth and eventual systemic colonization. Understanding the dynamics of crown infection is particularly important for developing targeted disease management strategies, potentially influencing approaches to fungicide application and cultural practices.

Although zoospores are the primary dispersal and infection propagules of *P. capsici* in the field, we initiated infections with standardized mycelial amounts. This choice provided (i) a synchronized and reproducible infection starting across hosts with distinct surface chemistries, (ii) sufficient, clean pathogen RNA from the crown interface for comparative transcriptomics, and (iii) reduced variability due to zoospore encystment and germination kinetics. Under our conditions, mycelial inoculation consistently established crown colonization across species (Fig. 1a-d), enabling uniform sampling at 24 hpi. We therefore interpret our RNA-seq data as capturing host-conditioned crown-associated hyphae; we do not claim that they substitute for all aspects of zoospore-initiated infection in the field. Future work will explicitly compare zoospore-versus mycelium-initiated infections to dissect any inoculum-type effects on early transcriptional programs. These profiles primarily reflect hyphae exposed to host-derived cues at the infection interface, not isolated intracellular hyphae. This design allows for clean pathogen RNA recovery and host-matched comparisons, but it introduces an anatomical bias. Notably, our microscopy results revealed that at 24 hpi, cucurbits and tomatoes already exhibited abundant intercellular growth in stems 1 cm above the inoculation site (Fig. 1f–i; Fig. S1), whereas chili peppers did not, even upon later inspection (Fig. S2). Accordingly, we view our RNA-seq data as crown-interface, host-conditioned states that likely correlate with but do not fully encompass the transcriptional programs of invasive hyphae within each host. This caveat is especially relevant for pepper, where limited tissue ingress at 24–48 hpi suggests that external hyphae may dominate the sampled signal. Future experiments using tissue-enriched sampling will refine the in-planta compartment-specific programs.

### 3.3 Metabolic reprogramming and nutrient acquisition

The ability of *P. capsici* to reconfigure its metabolic networks in response to host-specific conditions emerged as another central theme of our work. In tomato, the upregulation of glycolytic and fatty acid metabolism pathways suggests rapid mobilization of energy and biosynthetic precursors to fuel aggressive colonization, a scenario commonly observed in necrotrophic interactions (Kwaik and Bumann 2015; Petrasch et al. 2019; Williams and Lorenz 2020). In cucurbits, however, our transcriptomic data revealed a pronounced enrichment of pathways related to carbohydrate metabolism, galactose utilization, and glycerolipid metabolism. These alterations may allow *P. capsici* to exploit the nutrient-rich vascular environment of cucumber and melon, thereby supporting extensive intracellular colonization without triggering immediate cell death. Such a strategy resembles the concept of “stealth colonization”, wherein the pathogen minimizes damage to host tissues to secure a longer-term nutritional niche (Petrasch et al. 2019; Williams and Lorenz 2020).

In chili pepper (CM334, a genotype known to be partially resistant), the induction of metabolic pathways associated with cell wall modification and nitrogen compound metabolism might reflect adaptive responses to overcome the host’s defensive mechanisms. It is likely that *P. capsici* attempts to modify its own membranes and metabolic outputs to adapt better to a nutrient-scarce, hostile environment imposed by robust host immunity. Such reprogramming may help to adapt to a hostile microenvironment (Luo et al. 2021). This metabolic flexibility is emerging as a key determinant of pathogen virulence and host adaptation, as has also been observed in other oomycetes (Ah-Fong et al. 2017). The divergent metabolic signatures among hosts underscore the plasticity of *P. capsici* physiology and its capacity to fine-tune nutrient acquisition strategies based on host-derived signals. This versatility in metabolic reprogramming has been documented in other soil-borne pathogens and is increasingly recognized as a key determinant of virulence and host adaptation (Petrasch et al. 2019).

### 3.4 Effector Dynamics and Host-Specific Virulence

Effector proteins are key molecular tools employed by several pathogens to manipulate host immunity and promote colonization. Oomycete effectors are broadly categorized into cytoplasmic effectors and apoplastic effectors, depending on their localization and function (Bozkurt et al. 2011). Our transcriptomic analysis revealed that *P. capsici* effector expression is tightly regulated in a host-dependent manner. Among effector proteins, RxLR effectors are of particular interest because of their ability to translocate into host cells and suppress immune responses by targeting host signaling pathways (Wang et al. 2023; Whisson et al. 2007). Our data show that while many RxLR effectors are repressed, a subset is induced selectively depending on the host species. For example, homologs of PcAvh328 (Pc123179, Pc128001, and Pc552887), which have been shown to interfere with host defense signaling, displayed differential expression patterns, with at least one homolog being induced in chili pepper. Such induction in a typically resistant host may represent a targeted effort by the pathogen to counteract strong basal defenses. Two previously uncharacterized RxLR effectors (Pc503811 and Pc18476) were selected for functional analysis via dsRNA-mediated silencing. Pc503811, which is induced specifically during infection in chili pepper and tomato, and Pc18476, which is expressed in cucumber, melon, and tomato infections, play distinct host-dependent roles. Silencing of Pc503811 resulted in a modest (18%) reduction in pathogen growth on detached cucumber leaves, whereas a substantial 76% growth reduction was observed on detached tomato leaves. During stem infection of cucumber plantlets, silencing Pc503811 had no significant effect on pathogen growth. However, in melon stems, although pathogen growth remained comparable to that of the control, necrosis development was prevented. Notably, in chili pepper stems, pathogen growth was notably reduced, and in tomato stems, both pathogen growth reduction and necrosis prevention were observed. These results suggest that the function of Pc503811 may be particularly relevant in interactions with solanaceous hosts. The effector Pc18476, which is induced in all susceptible hosts, also has host- and tissue-dependent effects when silenced via dsRNA. In detached cucumber leaves, Pc18476 dsRNA treatment resulted in a substantial 78% reduction in pathogen growth; however, this treatment had no detectable effect on pathogen growth during infection of cucumber plantlet stems. In melon stems, pathogen growth was like in controls, yet necrosis development was prevented by dsRNA treatment. In chili pepper stems, pathogen growth was reduced by 54%, whereas in tomato stems, growth was modestly reduced by 18%, but importantly, necrosis was completely prevented.

PcAvh16 homologs (Pc14944, Pc14948, and Pc533084), which are implicated in the suppression of programmed cell death (Wang et al. 2011), are differentially regulated, suggesting that *P. capsici* tailors its effector deployment to prolong its biotrophic phase, particularly in hosts with rapid necrotic responses like tomato. In contrast, the consistent downregulation of effectors such as PcAvh166 across hosts may indicate roles confined to early infection events or environmental conditions not captured at our 24 hpi sampling. Moreover, our observations of elicitins (INF1 and INF2B) and Nep1-like proteins (NLPs) underscore the complexity of effector regulation. Canonical elicitins are well known from *P. infestans* to induce hypersensitive responses (Derevnina et al. 2016; Du et al. 2017; Huitema et al. 2005); their repression in cucurbits could be interpreted as deliberate suppression to avoid triggering host defenses prematurely. In contrast, the unique induction of PcNLP14 in chili pepper may represent a specialized mechanism to manipulate host cell death pathways and modulate immune responses (Chen et al. 2018).

The NLP effector Pc9358, which is specifically induced during infections of chili pepper, exhibited host- and tissue-dependent phenotypes when silenced by dsRNA. On detached cucumber leaves, Pc9358 silencing caused a 65% reduction in pathogen growth, although no effect was observed on cucumber plantlet stems. In melon stems, pathogen growth was unaffected by dsRNA treatment, but the induction of necrosis was completely prevented. In chili pepper and tomato stems, Pc9358 silencing reduced pathogen growth by 32% and 34%, respectively, and prevented necrosis development in both hosts. Interestingly, this was the only effective silencing treatment that successfully prevented necrosis in chili pepper stems.

We also selected Pc533030, encoding the conserved cell cycle regulator SDA1, as a control for evaluating the efficacy of dsRNA treatments on pathogen growth. This gene was non-differentially expressed in our transcriptomic datasets but was previously shown to be essential for sporangial morphology, mycelial growth, and virulence during susceptible chili pepper leaf infections (Zhu et al. 2016). Treatment of *P. capsici* with PcSDA1 dsRNA resulted in a 76% reduction in pathogen growth on detached cucumber leaves; however, pathogen growth unexpectedly increased by 10% in tomato leaves. In plantlet stems, PcSDA1 dsRNA treatment caused severe and aggressive necrosis in chili peppers and tomatoes, surpassing control infections and leading to complete tissue necrosis. No changes were observed in the cucumber stems, whereas necrosis was prevented in the melon stems.

The growth reduction observed in cucumber leaves is consistent with previous findings that SDA1 is critical for hyphal development and virulence. However, the increased growth in tomato leaves and exacerbated necrosis in tomato and chili stems are unexpected and may reflect a compensatory or stress-induced hypervirulence response when the cell cycle is disrupted (Goncheva et al. 2019). This raises the possibility that interference with essential cellular functions in *P. capsici* might dysregulate effector release or stress signaling, enhancing host tissue damage in some contexts (Goncheva et al. 2019). Conversely, the absence of necrosis in melon stems might indicate a differential host response to weakened or developmentally impaired pathogen tissue. Altogether, these findings illustrate the complexity of targeting conserved genes and highlight that silencing even core regulators like SDA1 can lead to variable outcomes depending on the infection environment, host genotype, or tissue type.

Recent reviews have described that the spatial and temporal regulation of effectors is crucial for pathogen success (Wang et al. 2023). Our study extends these findings by demonstrating that the expression profiles of effectors are not only host-specific but may also be influenced by the infection niche, in our case, the crown region comprising the stem base and roots. This localized deployment of effectors may reflect an adaptation to the unique defense barriers and nutrient landscapes encountered at the soil-plant interface, where rapid colonization of vascular tissues can be pivotal for systemic infection.

## 4 Conclusions

We found that *Phytophthora capsici* has developed an adaptive strategy to colonize diverse plant species. In tomato, rapid necrosis limits pathogen spread, yet aggressive metabolic reprogramming supports early infection. On the other hand, the slow necrosis induction in cucumber and melon coupled with extensive intercellular colonization points to a tolerance interaction that may facilitate prolonged pathogen proliferation. In the partially resistant chili pepper CM334, robust defense mechanisms inhibit intercellular colonization despite transcriptional changes in the pathogen. Our comprehensive pathogen-centered transcriptomic analysis revealed that differential effector deployment, metabolic reprogramming, and nutrient acquisition strategies are key components of *P. capsici* adaptive virulence. This study highlights the value of examining the transcriptional dynamics of pathogens, an approach often overlooked in favor of host-focused analyses. By focusing on crown infections, this work expands our understanding of soil-borne infection dynamics, a perspective that has been underrepresented in studies that predominantly target foliar diseases.

## 5 Materials and methods

### 5.1 Strains, media and culture conditions

The *Phytophthora capsici* strain used in this study was D3 mating type A1 (Pons-Hernández et al. 2020; Sevillano-Serrano et al. 2024). *P. capsici* D3 was routinely grown on V8 agar in the dark at 24°C.

### 5.2 Infection assay

To infect seedlings with *P. capsici* D3, 0.01 g of mycelium from 5-day-old cultures was harvested with tweezers and deposited in the crown of 5-8 weeks-old seedlings. Plants used as hosts in the present study were *Capsicum annuum* (Criollo de Morelos 334), *Cucumis melo*, *Cucumis sativus*, and *Solanum lycopersicum*. As a non-plant infection control, the same mycelium source was cultivated on V8 agar plates. At 24 hours post-inoculation, the accessible mycelium on the infected plant surface was recovered with tweezers for subsequent RNA extraction. To quantify necrosis at the crown, the plants were inspected at fixed intervals (every 6 h from 6-144 hpi). At each timepoint, a 50 mm segment encompassing the stem base and upper root segment centered on the inoculation site was excised and processed as follows to visualize dead cells: tissues were cleared overnight in acetic acid:ethanol (1:3, v/v) and then transferred to acetic acid:ethanol:glycerol (1:5:1, v/v/v) for 3 h. Cleared samples were stained for 3 h in 0.01% (w/v) trypan blue and stored in 60% glycerol until examination. Stained tissues were imaged at constant magnification via a bright-field microscope. Necrosis onset (event) was defined a priori as the earliest timepoint at which a contiguous trypan blue-positive band of ≥2.0 mm in longitudinal extent was present at the inoculation zone. Quantification was performed in ImageJ by RGB channel separation and intensity-thresholding of the blue channel within a fixed ROI (50 mm length × full stem width). For each host and time, 3 plants were scored across 3 independent experiments.

### 5.3 Multiphoton microscopy

Stems and roots were dissected from 1 cm from the crown inoculation site. The samples were fixed in 4% paraformaldehyde and incubated for 2 h. Subsequently, the samples were washed with 0.16 M phosphate buffer at pH 7.2, embedded in Sakura Tissue Tek, and frozen at -20°C. The samples were sectioned in a cryotome to a width of 100 μm. The tissue sections were stained with propidium iodine (IP) at a final concentration of 0.0003%. To stain the cell wall, the samples were treated with 0.01% Solophenyl. The stained samples were visualized via an LSM 880-NLO Axio Imager Z2 microscope (Zeiss, Oberkochen, Germany) coupled to a Ti:Sapphire infrared laser (Chameleon Vision II, COHERENT, Santa Clara, CA, USA). The excitation laser was set at 543 nm for propidium iodine and 458 nm for the solophenyl. The emission ranged from 548 to 684 for IP and 502 to 558 nm for the solophenyl. All the micrographs were captured in CZI format at 1131 × 1131 pixels and RGB color. The spectral graphs were analyzed via SigmaPlot 12 (Systat Software, Inc.).

### 5.4 RNA extraction and Illumina sequencing

Total RNA was extracted via the Quick-RNA Plant Miniprep Kit (Zymo Research, CA, USA) according to the manufacturer’s protocol, omitting the Zymo-Spin III-HRC column in the final step. Mycelia of 10 infected plants were pooled per extraction, and 0.1 g of mycelia was used for each extraction; mycelia were harvested at 12:00 (±1 hr), immediately flash-frozen in liquid nitrogen, and processed without thawing. The RNA quality was measured with a Bioanalyzer, and samples with a RIN > 5 were used for sequencing. RNA-Seq libraries (15 libraries in total, representing five conditions with three biological replicates per condition) were prepared by selectively isolating mRNA via poly-T oligo-attached magnetic beads. Libraries were sequenced at Novogene (Sacramento, CA, USA) via the Illumina NovaSeq 6000 platform (Hayward, CA, USA), generating paired-end reads (2 × 150 bp) with an average yield of 50 million reads per library. The raw sequencing data have been deposited in the NCBI database under BioProject accession number PRJNA1207595.

### 5.5 Mapping and statistical analysis

Read trimming and adaptor removal were performed using BBDuk with the following parameters: `forcetrimleft=20` and `trimq=20` (https://sourceforge.net/projects/bbmap/). Mapping and quantification were conducted with Kallisto via the *P. capsici* LT1534 v11.0 transcriptome (available at MycoCosm: mycocosm.jgi.doe.gov) (Bray et al. 2016; Grigoriev et al. 2014; Lamour et al. 2012a), employing bias correction and 100 bootstrap replicates. Approximately 77% of the reads were pseudoaligned to the transcriptome. Differential expression analysis was carried out via DESeq2, as described previously (Love et al. 2014; Tanja 2023). *P. capsici* grown on V8 agar served as the control condition. Transcripts with an adjusted p value (p value) ≤ 0.05 and a log2FoldChange > 1 or < -1 were considered differentially expressed genes (DEGs). Co-expression analysis was executed with Clust 1.18, using the parameters `–no optimization` (Abu-Jamous and Kelly 2018).

### 5.6 Functional enrichment

To identify gene ontology (GO) terms and enriched KEGG pathways, the reference transcriptome was reannotated via eggNOG mapper v2.1.12 (Huerta-Cepas et al. 2019) and merged with existing MycoCosm annotations. Effector proteins, including RXLRs, CRNs, Elicitins, and Nep1-like proteins (NLPs), were retrieved from the literature and supplemented by identifying additional candidates using EffectorP v3.0. Additional secreted proteins were predicted via SignalP v6 (Sperschneider and Dodds 2022; Teufel et al. 2022). Functional enrichment analysis was performed with clusterProfiler v4.12.6, employing an over-representation analysis (ORA) approach and applying a *p* value cutoff of ≤0.05 (Wu et al. 2021). Effectors were annotated against various databases with BLAST using an e-value < 1×10^-30^.

### 5.7 Functional validation by RNAi

Functional validation by gene silencing of three effector genes was conducted by designing RNAi sequences and their respective primers using SIREN v0.1.5 (https://github.com/pablovargasmejia/SIREN), and the T7 promoter was added to each primer (Table S1). To avoid off-target effects on the pathogen and host plants, an off-target database was constructed with the cDNA sequences of *P. capsici* and the four host plants with the --targets option in SIREN v0.1.5. Designed RNAi sequences were transcribed *in vitro* from *P*. *capsici* cDNA fused with the T7 promoter using HighYield T7 RNAi Kit at 37°C for 4 h (Jena Bioscience, Germany). RNAi experiments were performed as follows: 1 µg of synthetic dsRNA was incubated with 0.1 g of *P. capsici* mycelium for 2 h on ice to facilitate cellular uptake. Then, 0.01 g of treated mycelium was used as inoculum for virulence assays on adult plant leaves or plantlet stems. As a negative control for silencing, *P. capsici* mycelium was incubated as described with T7 reaction components that excluded RNA production. To screen the disease phenotype of the silenced genes, inoculated leaves and stems were first cleared in an acetic acid:ethanol (1:3, v/v) solution overnight and then in an acetic acid:ethanol:glycerol (1:5:1, v/v/v) solution for 3 h. The samples were subsequently incubated for 3 h in a staining solution of 0.01% (w/v) trypan blue and then stored in 60% glycerol until examination.

The reduction in target gene transcript levels following dsRNA treatment was quantified via RT-qPCR using the comparative ΔΔC_t_ method. The elongation factor 1 gene served as an endogenous gene. The oligonucleotide sequences are provided in Table S2. For each assay, three biological replicates were sampled at 6 hours post-inoculation (hpi) from either dsRNA-treated or control *P. capsici* mycelial inoculum. Total RNA was extracted as described previously, and 1 µg of RNA was reverse transcribed using the Jena SCRIPT cDNA Synthesis Kit (Jena Bioscience, Germany). Quantitative PCR was performed on an Applied Biosystems StepOne Real-Time PCR System (Thermo Fisher Scientific, USA), employing the qPCR GreenMaster highROX from Jena. Reaction conditions and cycling parameters followed the manufacturer’s guidelines.

## Declaration of competing interest

The authors declare no conflict of interest.

## Availability of data and materials

The raw reads generated during the current study are publicly available at NCBI under the PRJNA1207595 accession number. Materials are available from the corresponding author upon reasonable request. https://dataview.ncbi.nlm.nih.gov/object/PRJNA1207595?reviewer=3459l133boj3j0q7vu5nrpi1tp

## Acknowledgments

This work forms part of the requirements for P.V.M. to obtain a Doctoral degree from the Posgrado en Ciencias Biológicas program at Universidad Nacional Autónoma de México (UNAM). P.V.M. receives a doctoral scholarship from Secretaria de Ciencia, Humanidades, Tecnología e Innovación (SECIHTI). F.U.R.R. receives a posdoctoral fellowship from “Estancias Posdoctorales por México 2022” supported by SECIHTI. Funding was provided by PAPIIT-DGAPA-UNAM (project IN218024) and SECIHTI (project A1-S-34759) to JV-A.

## Notes

### Competing Interest Statement

The authors have declared no competing interest.

### Summary of Updates

Updated version of the manuscript. New discussion section, improved methods.

