## Supplementary material for "Adaptive transcriptional strategies underpin the host-specific virulence of the generalist oomycete *Phytophthora capsici* during early crown infection": Sup material

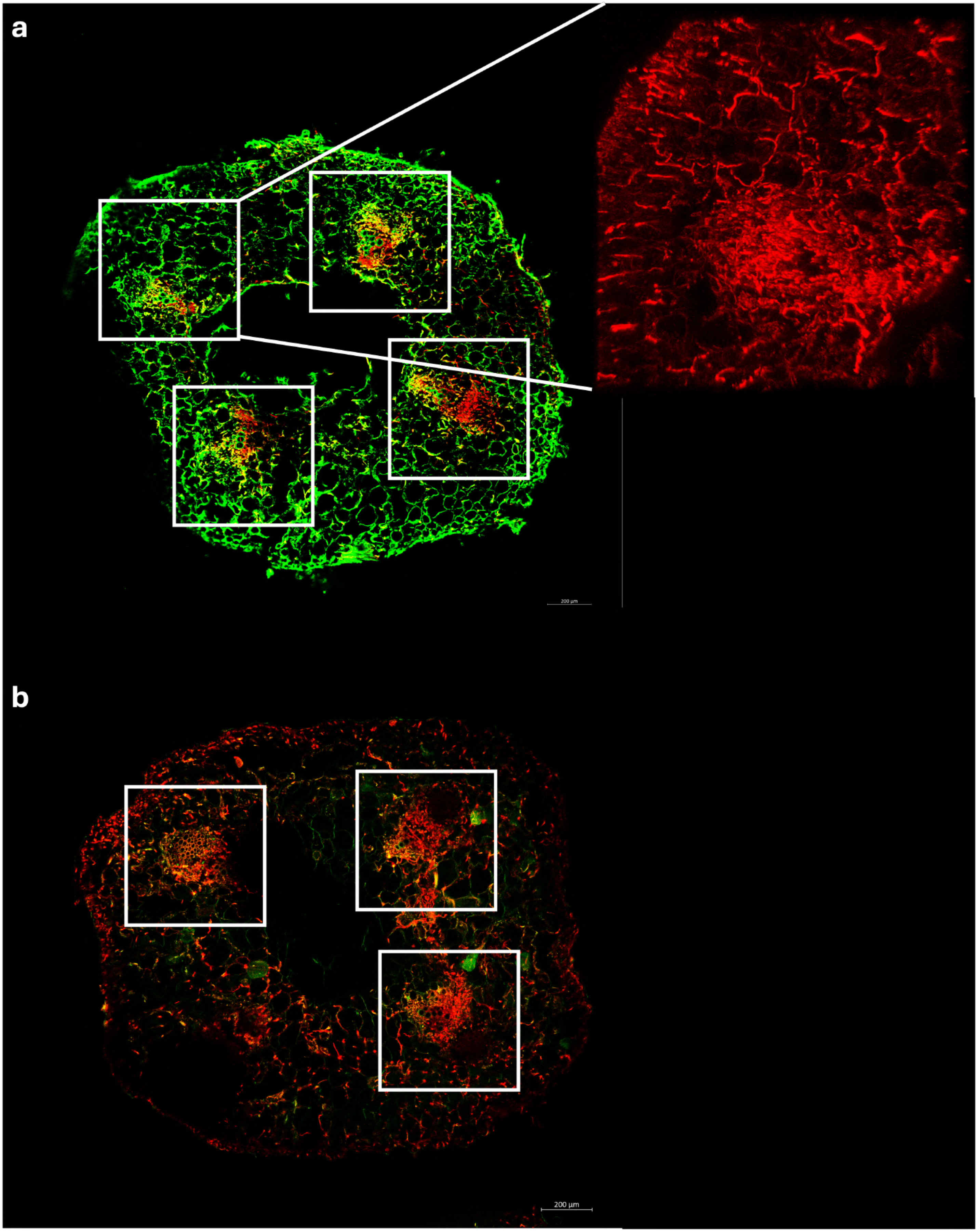


**Figure S1:** *Phytophthora* intracellular growth preference for vascular tissue in melon and cucumber. **a** Stem section of a melon-infected plant at 24 hpi with magnification of a section of vascular tissue. **b** Stem section of a cucumber plant at 24 hpi. Solophenyl (green channel) stains the plant cell wall, and propidium iodine (red channel) stans *Phytophthora* mycelium. White cubes highlight the vasculature.

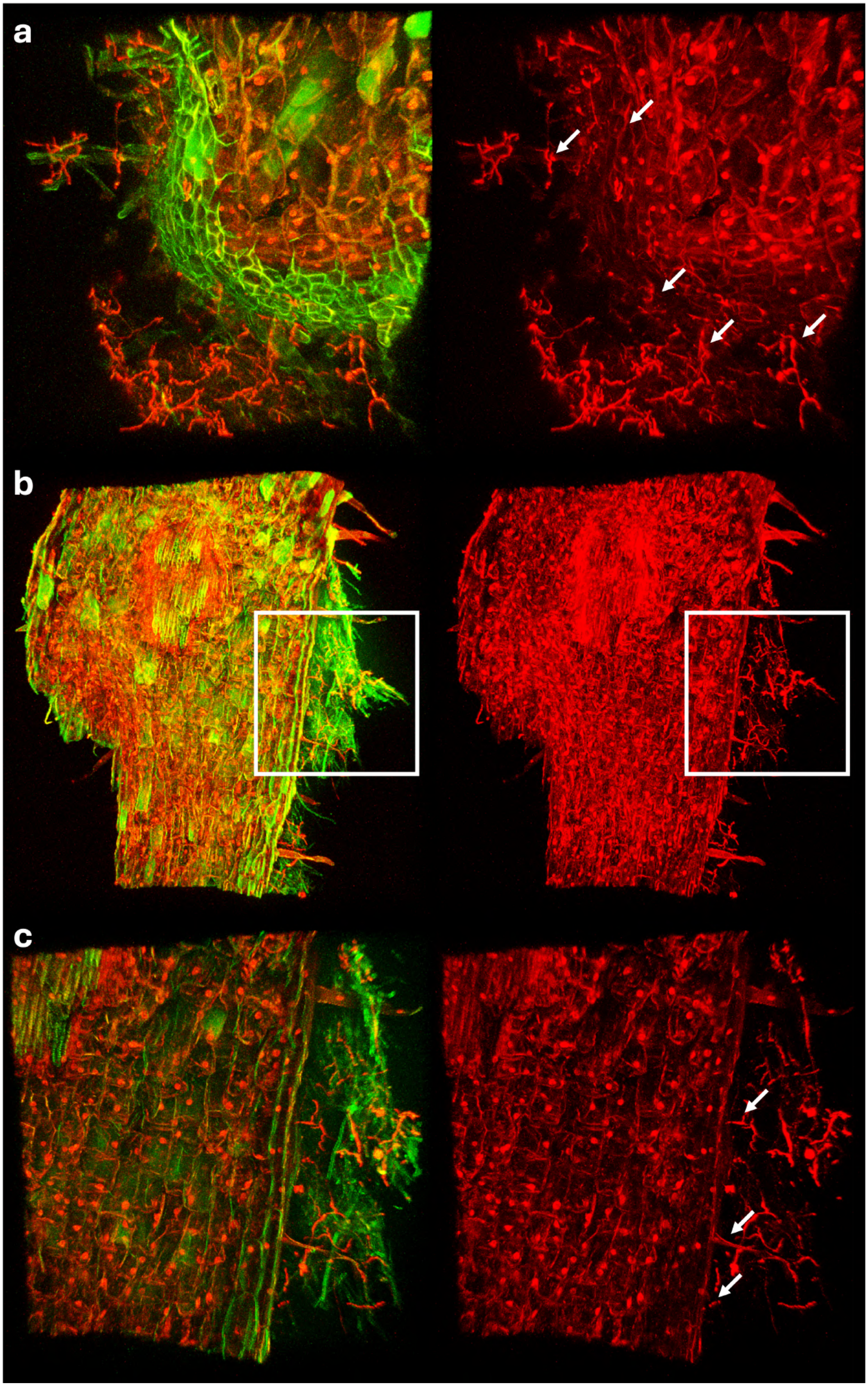
a
b
c

**Figure S2:** *Phytophthora* cannot penetrate CM334 pepper stems. **a** Inoculated crown section of pepper at 24 hpi. **b** Inoculated pepper stem section at 48 hpi. **c** Magnification of the inoculation site at 48 hr (as marked in panel b). Solophenyl (green channel) stains the plant cell wall, and propidium iodine (red channel) stains *Phytophthora* mycelium. The white cubes and arrows indicate the inoculation zone.

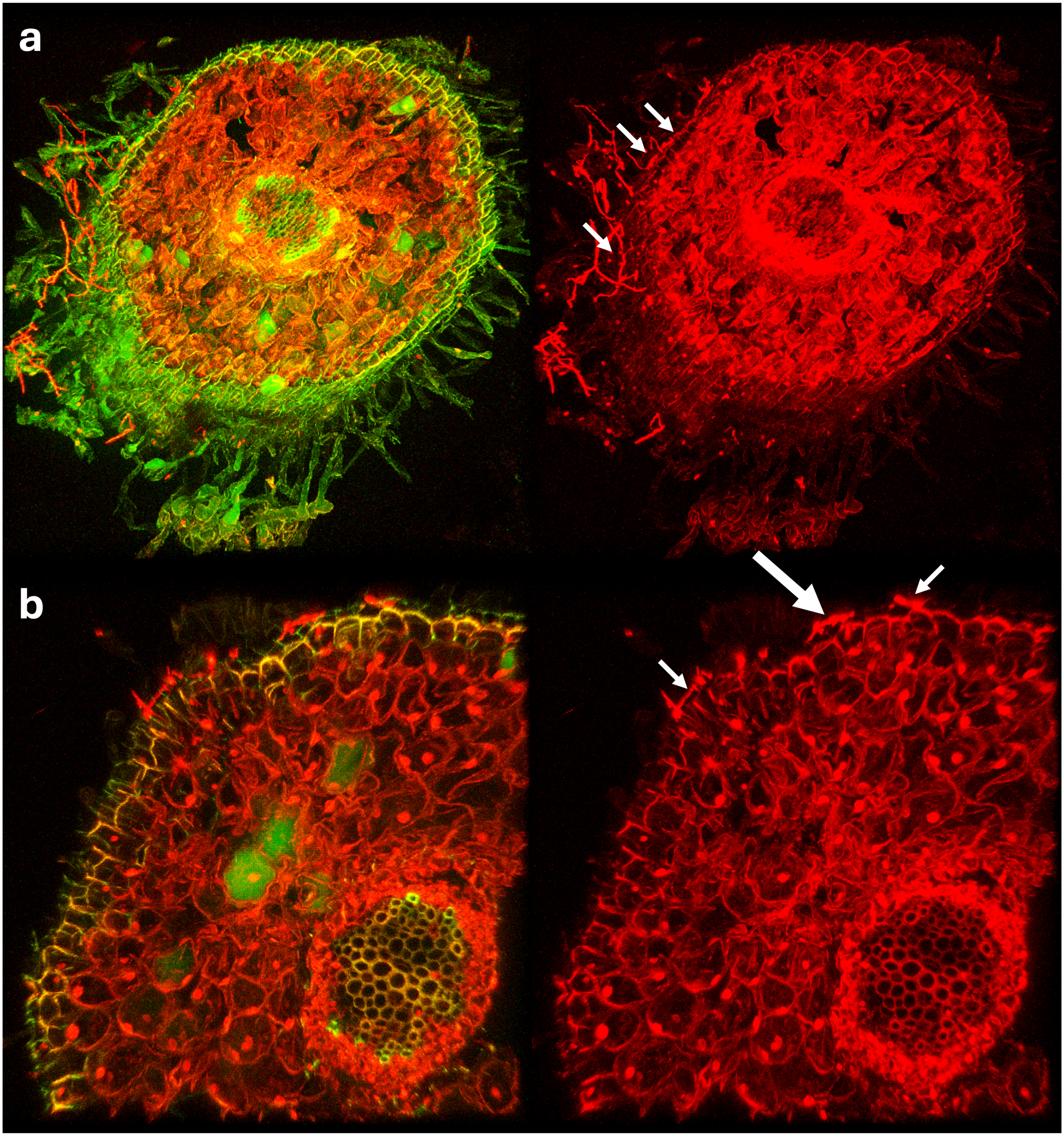


**Figure S3:** *Phytophthora* cannot penetrate CM334 pepper roots. **a** Inoculated root section of pepper at 24 hpi (10x). **b** Inoculated pepper root at 24 hpi with *P. capsici*-forming appressoria (large white arrow, 20x). Solophenyl (green channel)

stains the plant cell wall, and propidium iodine (red channel) stains *Phytophthora* mycelium. The white cubes and arrows indicate the inoculation zone.

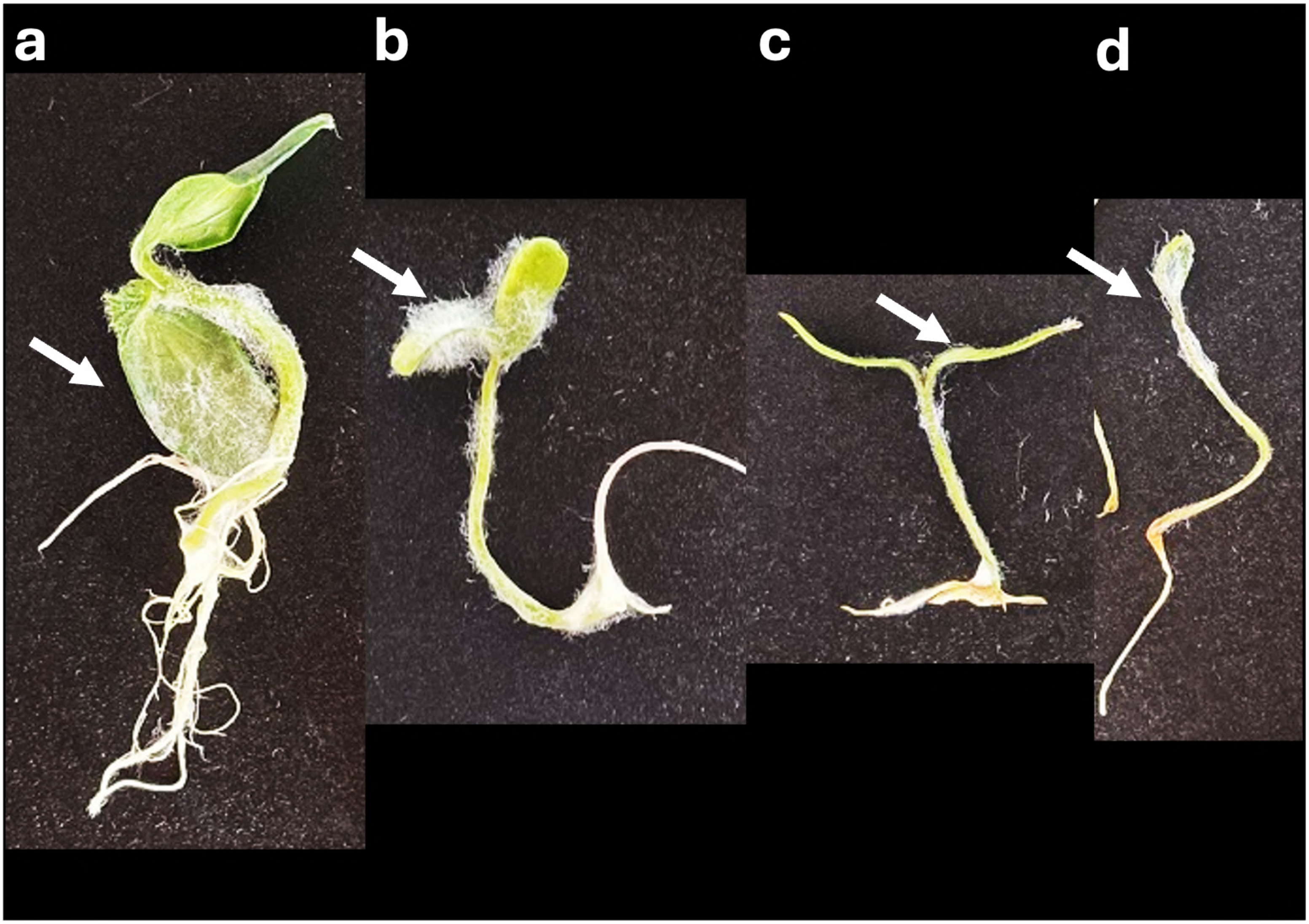


**Figure S4:** *Phytophthora* preference for aerial tissue. **a** Crown inoculated cucumber at 120 hpi. **b** Crown inoculated melon at 120 hpi. **c** Crown-inoculated pepper at 144 hpi d Crown-inoculated tomato at 96 hpi. The white arrows highlight mycelia on aerial tissue.

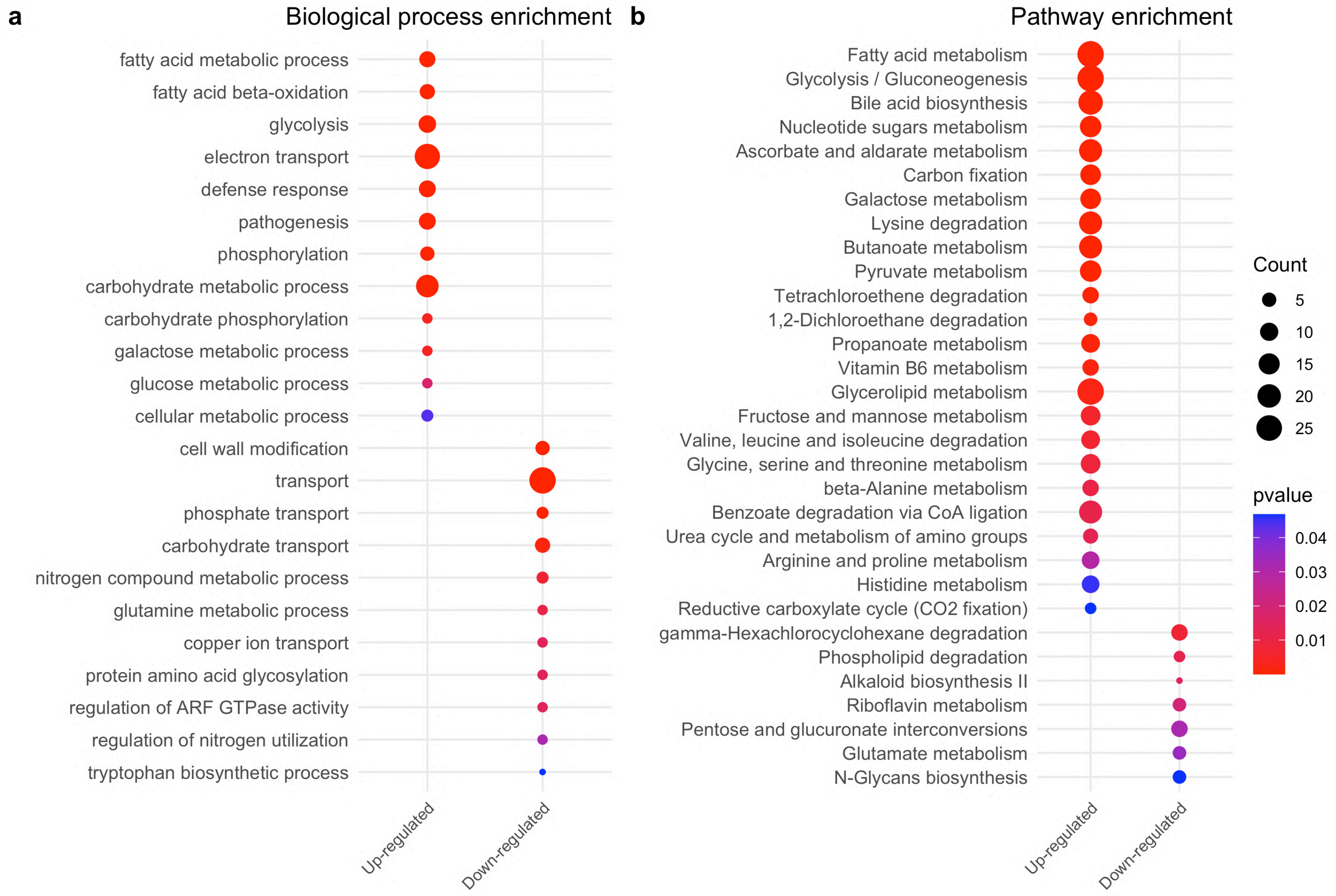


**Figure S5:** Core infection transcriptome enrichment. **a** Biological process enrichment (10x). **b** KEGG pathway enrichment.

**Table S1:** Primers and PCR conditions for dsRNA production.

| Primers | | | | PCR conditions first program | | | | PCR conditions second program | | | |
|---|---|---|---|---|---|---|---|---|---|---|---|
| Name | Length | Sequence | Expected size | denaturation | anneling | extension | # cycles | denaturation | anneling | extension | # cycles |
| 9358T7F | 38 | **TAATACGACTCACTATAGGG**ATCTTGGCTGTCACTCCT | 307 | 98°C 10s | 60°C 20s | 72.5°C 20s | x24 | 98°C 10s | 73.5°C 20s | 72°C 20s | x35 |
| 9358T7R | 38 | CCAGGCCTTGTTGAGTTT**GGGATATCACTCAGCATAAT** | | | | | | | | | |
| 503811T7F | 38 | **TAATACGACTCACTATAGGG**AAGAAGAGCGCGGTATTG | 312 | 98°C 10s | 60°C 20s | 72.5°C 20s | x24 | 98°C 10s | 73.5°C 20s | 72°C 20s | x35 |
| 503811T7R | 38 | GAGCTTGGCCACATCATC**GGGATATCACTCAGCATAAT** | | | | | | | | | |
| 18476T7F | 37 | **TAATACGACTCACTATAGG**GATCAGGCAGCCAAGTTT | 334 | 98°C 10s | 60°C 20s | 72.5°C 20s | x24 | 98°C 10s | 73.5°C 20s | 72°C 20s | x35 |
| 18476T7R | 38 | TTCTGACACCTTACGCAC**GGGATATCACTCAGCATAAT** | | | | | | | | | |
| PcSDA1 T7F | 38 | **TAATACGACTCACTATAGGG**CCAAGTGTTATCCCGTGG | 254 | 98°C 10s | 60°C 20s | 72.5°C 20s | x24 | 98°C 10s | 73.5°C 20s | 72°C 20s | x35 |
| PcSDA1 T7R | 36 | TATAGCAACTCACGCA**GGGATATCACTCAGCATAAT** | | | | | | | | | |

**Table S2:** Primers for RT–qPCR validation of dsRNA-induced gene silencing.

| Primers | |
|---|---|
| Name | Sequence |
| 9358Fw | CCT TCA GCT CAC AGT GGG TA |
| 9358Rv | TTG CTC CCA CAT GAT GAG GT |
| 503811Fw | TGA CGA GGT GCT TAA GGT GT |
| 503811Rv | CTT GCT TCT CAC TGA GCG AC |
| 18476Fw | AAG AAA GCC AAC ACT GAC GC |
| 18476Rv | CGT TAG CCG CTA CCT TGA AC |
| PcSDA1Fw | TGG ATG GGA AGT TGC CTC AT |
| PcSDA1Rv | AAA GTC AAG AGG CGT CAG GA |
| EF1Fw | CTA AGG GCA CCC AGG ACT TC |
| EF1Rv | TCA CCC GAC TTC ACG AAC TT |
